# An intrinsic hierarchical, retinotopic organization of pulvinar connectivity in the human neonate

**DOI:** 10.1101/2024.07.30.605817

**Authors:** Vladislav Ayzenberg, Chenjie Song, Michael Arcaro

## Abstract

Thalamic connectivity is crucial for the development of the neocortex. The pulvinar nuclei are thought to be particularly important for visual development due to their involvement in various functions that emerge early in infancy. The development of these connections constrains the role the pulvinar plays in infant visual processing and the maturation of associated cortical networks. However, the extent to which pulvino-cortical pathways found in adults are established at birth remains largely unknown, limiting our understanding of how these thalamic connections may support infant vision. To address this gap, we examined the organization of pulvino-cortical connections in human neonates using probabilistic tractography analyses on diffusion imaging data. Our findings revealed the presence of white matter pathways between the pulvinar and each individual visual area at birth. These pathways exhibited specificity in their connectivity within the pulvinar, reflecting both intraareal retinotopic organization and the hierarchical organization across ventral, lateral, and dorsal visual cortical pathways. These connections could enable detailed processing of information across sensory space and communication along distinct processing pathways. Comparative analyses revealed that the large-scale organization of pulvino-cortical connectivity in neonates mirrored that of adults. However, connectivity with ventral visual cortex was less adult-like than the other cortical pathways, aligning with prior findings of protracted development associated with the visual recognition pathway. These results deepen our understanding of the developmental trajectory of thalamocortical connections and provide a framework for how subcortical structures may support early perceptual abilities and scaffold the development of cortex.

## Introduction

From face recognition to motion perception and visuospatial reasoning, human infants exhibit remarkable perceptual abilities within the first neonatal days (Gava et al., 2009; Johnson et al., 1991; Sai, 2005; Spelke et al., 1995; Turati et al., 2008). This early perceptual competence is striking, given that infant visual sensitivity and acuity are initially poor and exhibit a prolonged postnatal maturation (Kiorpes, 2016). In adults, visual behaviors are supported by a network of cortical areas distributed along ventral, lateral, and dorsal visual pathways (Haak & Beckmann, 2018; Kanwisher & Dilks, 2012; Pitcher & Ungerleider, 2021; Ungerleider & Haxby, 1994). While the basic architecture and response properties of early visual cortex are surprisingly mature in infants (Kiorpes, 2016; Wiesel & Hubel, 1974), many aspects of this neocortical circuitry, particularly in higher-order extrastriate cortex, remain immature and undergo refinement throughout postnatal development (Bourne & Rosa, 2006; Golarai et al., 2007; Sydnor et al., 2023).

This discrepancy between advanced perceptual abilities and immature cortical circuitry has led researchers to hypothesize that early infant visual behaviors are supported primarily by subcortical structures (Alcauter et al., 2014; Atkinson, 1983; Blumberg & Adolph, 2023; Johnson, 2005). The pulvinar, in particular, may be critical for early infant visual behavior (Arcaro et al., 2018; Bridge et al., 2016). In adults, the pulvinar engages in diverse visual functions, processing both low-level features such as local contrast and orientation (Bender, 1982) to high-level features such as whole objects and faces (Ngyuen et al. 2013; Arcaro et al. 2018), as well as visual motion (Dumbrava et al., 2001; Merabet et al., 1998). Importantly, the structure of the pulvinar (Homman-Ludiye et al., 2018; Kostovic & Vasung, 2009; Rakić & Sidman, 1969) including its internal subdivisions (Ogren & Rakic, 1981) and cortical connectivity (Ball et al., 2013; Counsell et al., 2007; Shatz & Rakic, 1981) begin to emerge during gestation. The pulvinar’s importance in early visual processing is further underscored by studies showing that damage to the pulvinar in infancy results in greater visual deficits than damage to corresponding cortical visual areas (Bourne & Morrone, 2017; Dubowitz et al., 1986). Moreover, the pulvinar may provide an alternate path for visual processing when primary visual cortex is damaged, preserving some visual functions in cortically blind individuals (Kinoshita et al., 2019; Leopold, 2012; Warner et al., 2015; but, see Ajina & Bridge, 2018). These findings suggest that maturation of the pulvinar constrains the development of a variety of visual behaviors.

Beyond its direct involvement in infant visual behavior, the pulvinar may be crucial in establishing visual cortical circuits. Thalamocortical connections have long been recognized as essential for the development of primary cortical fields (O’Leary et al., 2007), and recent evidence extends this importance to higher-order neocortical organization (Murakami et al., 2022). In mice, spontaneous thalamic activity helps drive the development of sensory regions (Aníbal-Martínez et al., 2024; Antón-Bolaños et al., 2019) and prenatal disruptions of thalamocortical connections alter the borders between visual areas (Chou et al., 2013) or prevents their development entirely (Antón-Bolaños et al., 2019). For higher-order visual circuits, the first connections between the pulvinar and extrastriate cortex emerge during primate gestation shortly after the lateral geniculate innervates primary visual cortex (Shatz & Rakic, 1981), setting the stage for the pulvinar to influence the formation of visual cortical circuits. Indeed, pulvinar connectivity has been shown to guide the initial development of area MT (Bourne, 2010; Mundinano et al., 2018; Warner et al., 2012). Recent research indicates an even broader role for the pulvinar in shaping visual cortical development. Thalamus connectivity in young children is a better predictor of large-scale cortical organization across development than local wiring rules (Park et al., 2024), highlighting the pulvinar’s potential influence on the overall architecture of the developing visual system.

Despite mounting evidence for the pulvinar’s importance in early cortical development and visual functioning, few studies have examined the organization of pulvino-cortical connectivity in human neonates. Previous work has shown that large-scale white matter connections between the pulvinar and cortex exist in human infants (Ball et al., 2013; Counsell et al., 2007), but have not examined the specificity or maturity of these connections. Here, we address this gap by examining how connectivity with the pulvinar varies across functionally distinct cortical pathways, and assessing to what extent these connections are adult-like at birth. We analyzed newborn diffusion tensor imaging (DTI) data from the developing human connectome project (dHCP) to provide a detailed analysis of the connectivity between the pulvinar and individual visual cortical areas across occipital, ventral-temporal, lateral-occipital, and dorsal-parietal visual cortices. By revealing the organization of pulvino-cortical connectivity in neonates, we aim to elucidate potential mechanisms underlying early perceptual abilities and identify principles supporting cortical development and early life plasticity.

## Results

### Pulvino-cortical anatomical tractography in neonates

We evaluated the anatomical organization of the pulvinar in neonatal humans using probabilistic tractography analyses on diffusion MRI. We assessed the extent to which white matter pathways between the pulvinar and 17 cortical visual areas (V1, V2, V3, hV4, VO1, VO2, PHC1, PHC2, LO1, LO2, MT, V3A, V3B, IPS0, IPS1, IPS2, IPS3) are present at birth. To determine the locations of individual cortical visual areas in neonates, we aligned an adult probabilistic atlas of retinotopic areas to each neonate’s high-resolution anatomical image (see Methods). Only ipsilateral (i.e., within hemisphere) connectivity was assessed.

### Identifying pulvino-cortical pathways

An initial probabilistic tractography analysis revealed the white matter thalamic radiation connecting the pulvinar with each cortical visual area. In each neonate, we identified white matter pathways linking individual cortical areas to the posterior thalamus (Figure 1A). The spatial locations of these pathways were consistent across neonates. Specifically, posterior occipital (e.g., V1; Figure 1A-B, 1^st^ row), ventral-temporal (e.g., VO1; Figure 1A-B, 2^nd^ row), lateral-occipital (e.g., MT; Figure 1A-B, 3^rd^ row), and dorsal-parietal (e.g., IPS3; Figure 1A-B, 4^th^ row) areas showed tracts connecting with the pulvinar through the posterior thalamic radiation (Figure 1B; Arcaro et al., 2015; Le Gros Clark & Boggon, 1935; Leh et al., 2008). The thalamic radiation for each cortical area entered the pulvinar laterally but varied along the dorsal-ventral axis (Figure 1A-B, 5^th^ row). Additionally, the thalamic radiation for posterior occipital and lateral occipital areas separated along the medial-lateral axis. These data demonstrate that the major white matter pathways linking the pulvinar and each visual area are present at birth.

**Figure 1.**
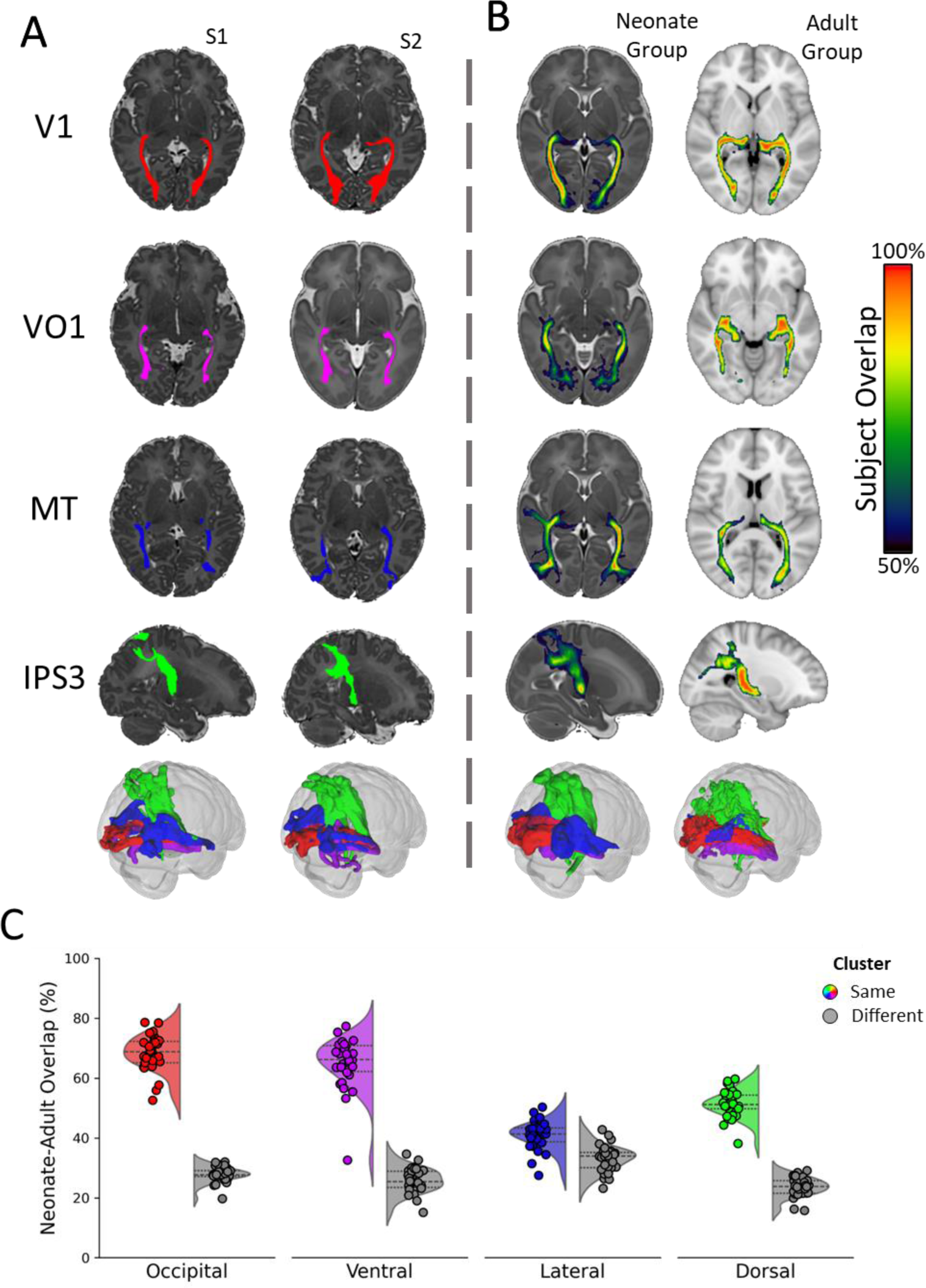
White matter pathways linking the pulvinar and visual cortex in neonates and adults. (A) White matter pathways between the pulvinar and occipital (V1), ventral (VO1), lateral (MT), and dorsal (IPS3) visual areas for two example neonates. (B) Group maps illustrating the degree of overlap for these pathways across individuals in neonates (left) and adults (right). (A-B; bottom) Three-dimensional glass brains illustrating the relative locations of white matter pathways between the pulvinar and V1 (red), VO1 (purple), MT (Blue), and IPS3 (Green) in neonates. For adults, white matter pathways reflect connectivity to each visual network as a whole. Tractography was performed ipsilaterally and combined across hemispheres for visualization purposes. (C) The degree of anatomical overlap between neonate and adult white matter pathways for occipital, ventral, lateral, and dorsal visual cortices. Colored distributions indicate the overlap between the same, corresponding, networks in neonates and adults, and gray distributions indicate the overlap between the neonate network and different, non-matching, networks in adults.

The white matter pathways interconnecting occipital, ventral, lateral, and dorsal cortical visual areas and the pulvinar identified in neonates were consistent with those observed in adults. A direct spatial comparison of tractography between neonates and adults (see Methods) revealed a main-effect of cluster similarity, such that there was greater overlap between tracts from the same cluster (e.g., mean overlap of neonate V1, V2, V3 with adult occipital cortex) than from different clusters (e.g., mean overlap of neonate V1, V2, V3 with adult lateral cortex; *F*(1,29) = 2350.64, *p* < .001, 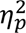 = 0.98; Figure 1C). Post-hoc comparisons (Holms-Bonferroni corrected) confirmed that there was adult-like specificity in the locations of white matter pathways for each visual cluster in neonates (Occipital: *d* = 7.97; Ventral: *d* = 7.71; Lateral: *d* = 1.54; Dorsal: *d* = 5.43; all *ps* < .001; Figure 1C). Similar specificity was observed when comparing individual neonate areas to adult clusters (Supplemental Figure 3). The pathway correspondence between neonates and adults for areas within the lateral (LO1, LO2, and MT) and dorsal (V3A and V3B) clusters exhibited smaller overlap values relative to occipital and ventral clusters. This difference likely stems from the higher degree of crossing fibers and tract curvature in these regions (Figure 1A-B, 5^th^ row; Supplemental Figure 3). Nevertheless, the specificity of lateral and dorsal tracts was still apparent in neonates. These data indicate the presence and specificity of the white matter pathways interconnecting visual cortex and the pulvinar in neonates comparable to adults.

### Cortical connectivity within the pulvinar

Having identified the white matter pathways interconnecting the visual cortex and the pulvinar in neonates, next we assessed which regions of the neonate pulvinar were anatomically interconnected with each cortical area. Probabilistic tractography for each cortical area was constrained to the thalamic radiation identified in the initial analysis.

Within the pulvinar, the highest tracking values for each area were confined to a focal region. The lateral-ventral pulvinar exhibited the highest tractography values for occipital (e.g., V1) and ventral cortical areas (e.g., VO1) (Figure 2, top two rows), while the dorsal-medial pulvinar had the highest tractography values for lateral (e.g., MT; Figure 2, 3^rd^ row) and dorsal cortical areas (e.g., IPS3; Figure 2; 4^th^ row). Thus, as in adulthood, visual cortical areas exhibit specificity in their connections within the pulvinar.

**Figure 2.**
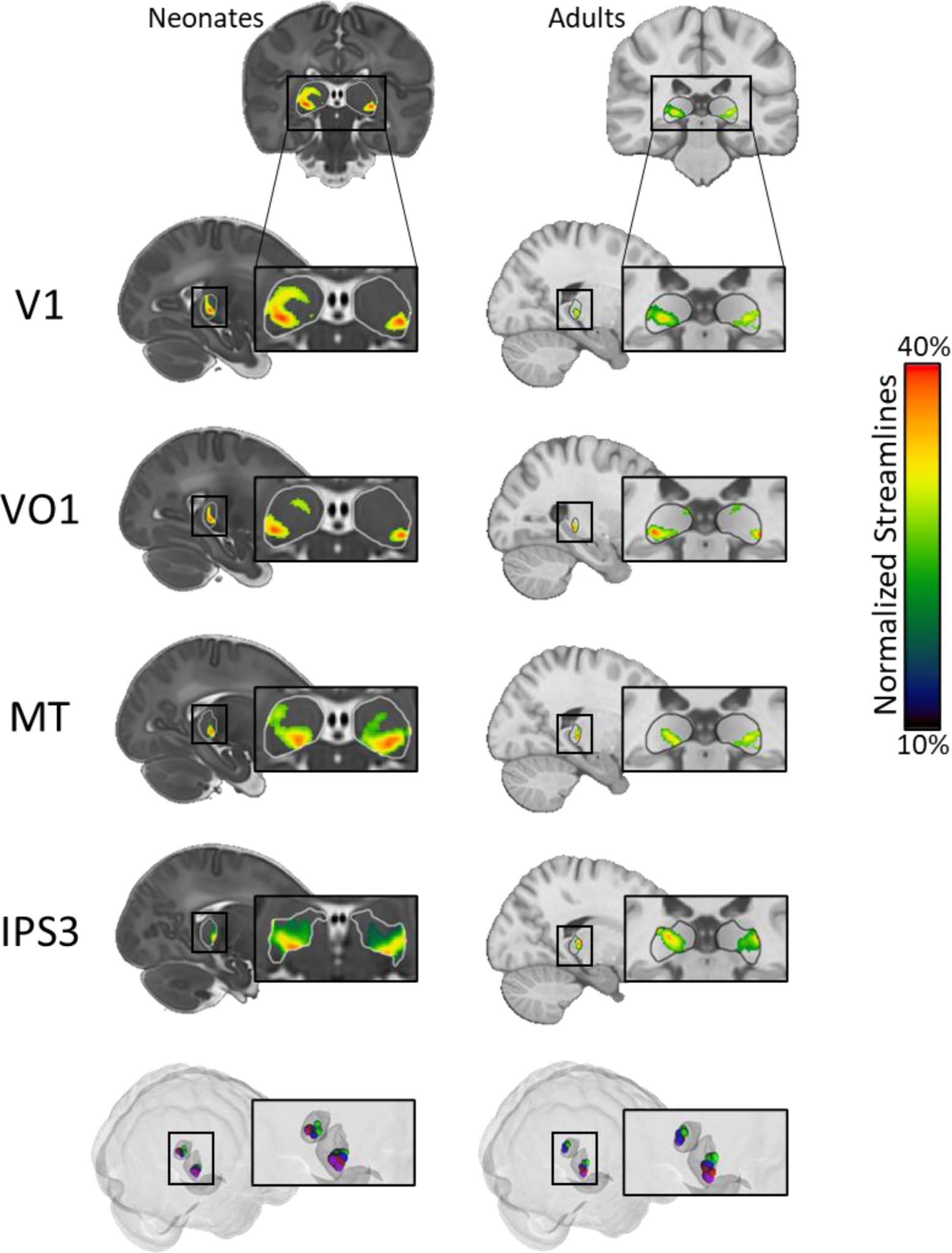
Localization of cortical connectivity within the pulvinar. Normalized peak streamline values within the pulvinar for the neonatal (left) and adult (right) group averages. All tractography was performed ipsilaterally and individual hemisphere data were combined for visualization. Three-dimensional glass brains (bottom) illustrate the relative locations of peak tracking values (top 10%) between the pulvinar and V1 (red), VO1 (purple), MT (Blue), and IPS3 (Green).

Next, we examined the organization of areal connectivity within the neonate pulvinar in relation to the visual pathway organization of cortical areas in adults. To this end, we computed pairwise correlations between the pulvinar tractography maps for each visual cortical area (Figure 2), creating a matrix depicting the similarity of connectivity patterns across all visual areas. Multi-dimensional scaling of this matrix revealed a systematic organization of cortical visual areas in neonates (Figure 3A). The resulting visualization showed two distinct progressions of cortical areas, both originating from occipital areas (V1-V3; red labeled areas). One progression followed the ventral cortical hierarchy (purple labeled areas), while the other encompassed both lateral (blue) and dorsal (green) cortical hierarchies. Interestingly, while connectivity between cortical visual areas is organized into three pathways (Kravitz et al., 2011; Kravitz et al., 2013; Pitcher & Ungerleider, 2021; Van Essen et al., 1992), anatomical connectivity with the pulvinar primarily reflected two pathways, with the lateral and dorsal cortical pathways not well differentiated. This organization was qualitatively similar to the structure observed in adults (Figure 3B; Arcaro et al. 2015), indicating that the broad topological organization of pulvino-cortical connectivity is already present at birth and broadly recapitulates the organization of cortical visual pathways.

**Figure 3.**
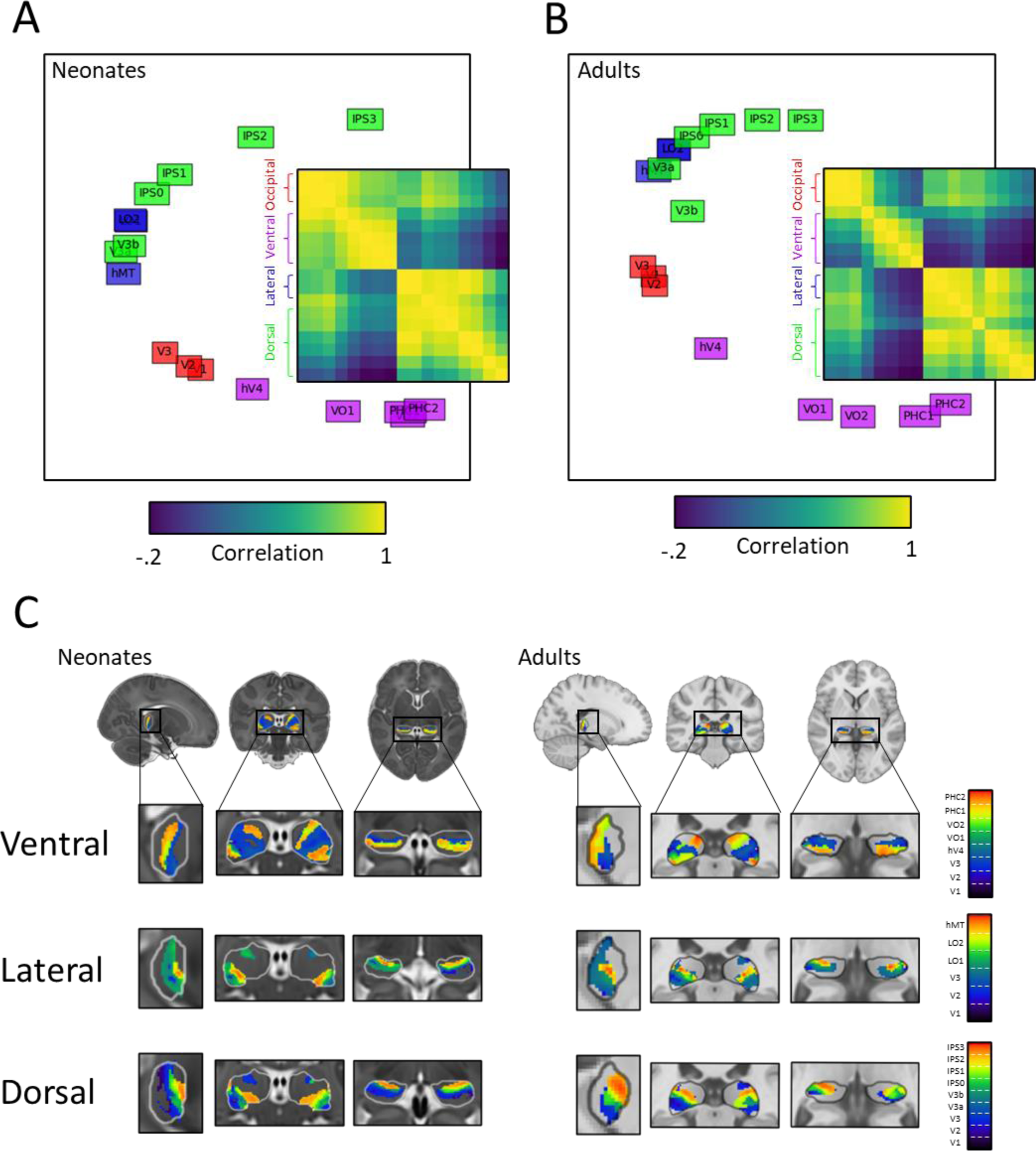
Connectivity patterns within the pulvinar. (A-B) Projection of first two principal components from multidimensional scaling of the dissimilarity between seed maps in the pulvinar with each visual area inside the pulvinar for (A) neonates and (B) adults. The inset shows the dissimilarity matrix comparing the correlations between connectivity maps in the pulvinar with each visual area. (C) Connectivity gradients for areas of the ventral, lateral, and dorsal visual pathways. Lower values (blue) correspond to the most posterior regions in each network’s hierarchy (occipital areas) and higher values (red) correspond to the most anterior regions in the hierarchy. Occipital areas are included in each figure gradient because they are the starting point of each cortical visual pathway.

In adults, these cortical pathways are hierarchically organized, originating in V1 at the posterior occipital pole and extend anteriorly along ventral, lateral, and dorsal cortex (Kravitz et al., 2011; Kravitz et al., 2013; Pitcher & Ungerleider, 2021; Van Essen et al., 1992). To examine whether the neonate pulvinar exhibits connectivity gradients that align with the hierarchical organization of ventral, lateral, and dorsal visual pathways, we assigned each pulvinar voxel a scalar value reflecting the visual area with the strongest connectivity to that voxel (see Methods). In the resulting gradient maps (Figure 3C), lower values (blue) correspond to the posterior occipital areas and higher values (red) correspond to the anterior areas within each cortical pathway. The gradient maps were similar for neonates and adults, with the starting point of each gradient localized to ventral-lateral portions of the pulvinar in both age groups, corresponding to early visual cortex. Connectivity across the ventral pathway revealed an anterior-to-posterior gradient, with the most anterior areas of the pathway localized to the posterior pulvinar (Figure 3C, 1^st^ row). In contrast, connectivity across both lateral and dorsal pathways exhibited a gradient moving from lateral-ventral to anterior-medial-dorsal portions of the pulvinar (Figure 3C, 2^nd^ and 3^rd^ row). Notably, the dorsal cortical pathway’s gradient map extended into more dorsal portions of the pulvinar compared to the lateral pathway map. These results demonstrate that visual cortical areas exhibit distinct connectivity patterns within the neonate pulvinar, similar to the hierarchical organization prominent in adults.

### Maturity of pulvino-cortical connectivity

Having characterized the organization of pulvino-cortical connectivity in neonates, we next assessed the maturity of these pathways by directly comparing the similarity of tractography between neonates and adults. Each neonate pulvino-cortical tractography map was correlated with each adult topography map (see Methods). These analyses revealed a main-effect of cluster similarity, such that neonates showed stronger correlations with adult tractography maps for cortical areas within the same cluster (e.g., neonate V1 to adult V1) than different clusters (e.g., neonate V1 to adult MT; Figure 4A) (*F*(1,29) = 808.43, *p* < .001, 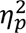 = .51). Post-hoc comparisons (Holms-Bonferroni corrected) confirmed that neonates exhibited such specificity for every cluster (Figure 4A; Occipital: *d* = 3.48; Ventral: *d* = 1.07; Lateral: *d* = 2.65; Dorsal: *d* = 5.24; *ps* < .001).

**Figure 4.**
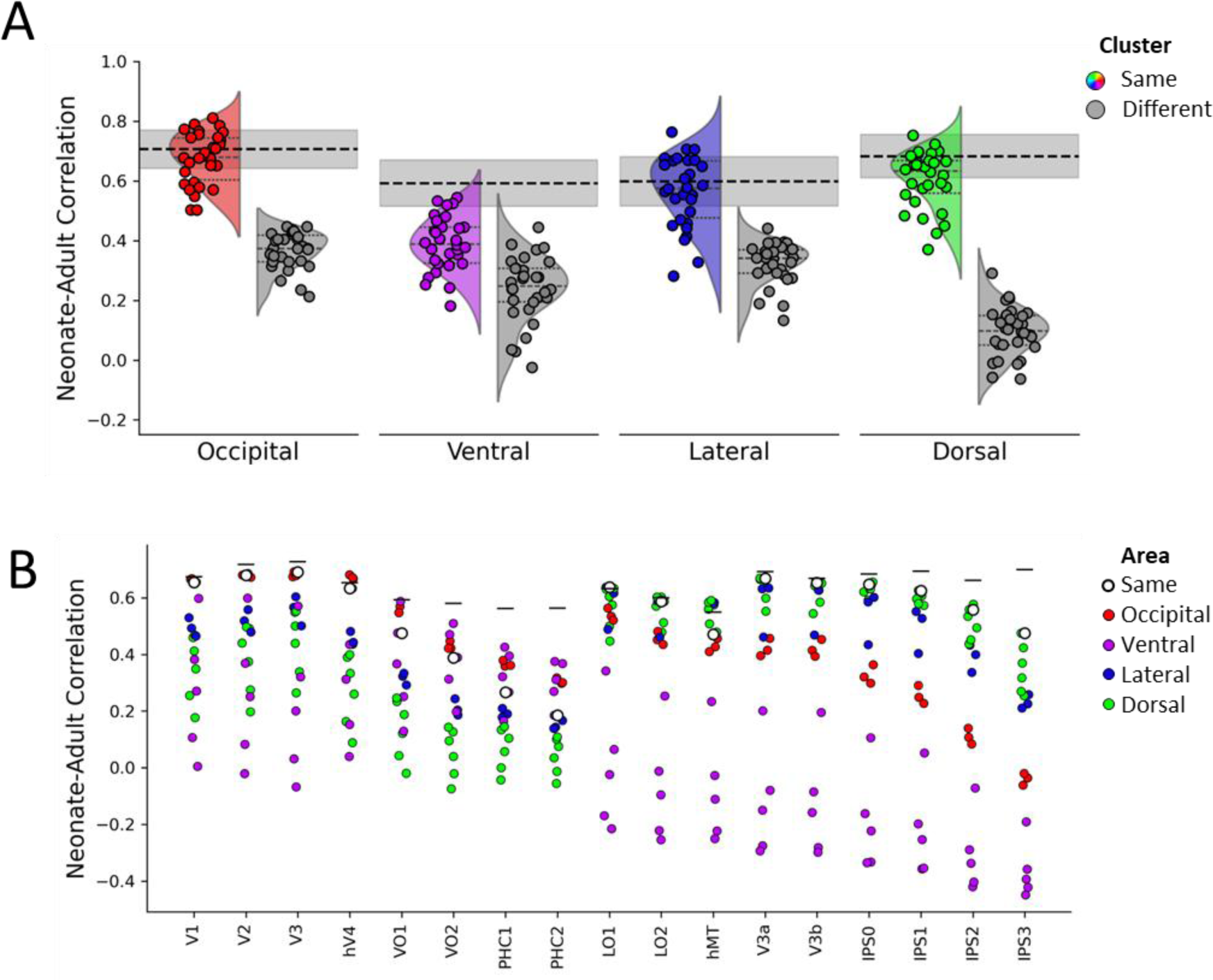
Comparisons of cortical tractography within the pulvinar between neonates and adults. (A) Cluster-level similarity between neonates and adults for occipital, ventral, lateral, and dorsal visual areas. Colored distributions show the correlation between anatomically clustered visual areas in neonates with the same cluster in adults. Gray distributions show the correlation with different clusters. The dotted black line with shading depicts the adult noise ceiling. (B) Area-level similarity between neonates and adults for all visual areas measured. White dots indicate the correlation with the corresponding area in adults and colored dots indicate the correlations with other visual areas. Visual areas are color coded by anatomical cluster. Black lines depict the adult noise ceiling for each region.

However, there were differences in the degree of the correspondence between neonate and adult maps across visual clusters. We measured each cluster’s overlap with the adult noise ceiling (Figure 4A; grey dashed line), reflecting the average correlation between individual adult participants and the mean of the remaining adults. Overlap was measured by the number of neonates whose correlations were greater than the lower-bound of the noise ceiling confidence intervals. Ventral maps showed less overlap with the adult noise ceiling (Ventral: 4/30 neonates) than the other visual clusters (Occipital: 21/30; Lateral: 21/30; Dorsal: 17/30). A comparison of the strength of each neonate correlation with adults (after normalizing by the noise ceiling), revealed a main-effect of cluster (*F*(3, 87) = 58.85, *p* < .001, 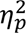 = .67). Specifically, post-hoc comparisons (Holms-Bonferroni corrected) revealed that correlations for the ventral areas were significantly lower than every other cluster (Occipital: *d* = 2.89; Lateral: *d* = 1.77; Dorsal: *d* = 2.17; *ps* < .001). Interestingly, occipital area correlations were also higher than lateral (*d* = 1.12, *p* < .001) and dorsal (*d* = 0.72, *p* = .004) clusters. There was no significant difference between lateral and dorsal clusters (*d* = 0.40, *p* = .082). These findings indicate that, at the area cluster level, the spatial specificity of the tractography for occipital, lateral, and dorsal cortices within the pulvinar are comparable to adults whereas pulvinar connectivity with the ventral cortical pathway is immature.

Finally, these differences in maturation of the pulvino-cortical tractography were apparent for individual cortical visual areas. For occipital, lateral, and dorsal cortices, we found that the correlation between corresponding areas in neonate and adults (e.g., neonate V1 with adult V1) was typically within the top three correlations of all area-to-area correlations (e.g., neonate V1 with adult V2; Figure 4B). The only exceptions in occipital, lateral, or dorsal cortices were LO2, which was the fourth highest correlation, and MT, which was the ninth highest correlation. Ventral areas, in contrast to areas of other clusters, did not show evidence of specificity (Figure 4B). These findings illustrate area specificity of neonate pulvino-cortical connections for occipital, lateral, and dorsal areas comparable to what is present in adults, but ventral areas exhibit more diffuse connectivity.

In addition to these maturation differences across visual pathways, the maturity of pulvino-cortical connectivity varied across the visual hierarchy. Correlations relative to the adult noise ceiling decreased as a function of distance from V1, with the strongest correlations occurring for the most posterior areas within a cortical visual pathway (Ventral: hV4; Lateral: LO1; Dorsal: V3a/IPS0) and relatively weaker correlations occurring for the most anterior areas (Ventral: PHC2; Lateral: MT; Dorsal: IPS3). The relative correlation strengths between neonates and adults for areas within ventral (V1 to PHC2), lateral (V1 to MT), and dorsal (V1 to IPS3) pathways was best fit by a two-feature polynomial (Ventral: *R^2^* = 0.54; Lateral: *R^2^* = 0.08; Dorsal: *R^2^* = 0.54), whose trajectory is relatively flat for posterior regions and then decreases moving anterior along each visual cortical pathway (Figure 5). Slope analysis for each pathway further revealed that ventral, lateral and dorsal pathways had a negative slope significantly different from 0 (*ps* < .001). A follow-up analysis revealed a significant difference in slope fits across regions, *F*(3, 87) = 19.17, *p* < .001, 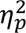 = 0.40, with post-hoc tests (Holms-Bonferroni corrected) showing that the fit across ventral areas had a significantly larger negative slope than the other two pathways (*ps* < .001). Thus, the maturity of pulvino-cortical connections varies across the visual hierarchy, with connectivity to higher-level areas being less mature than lower-level areas.

**Figure 5.**
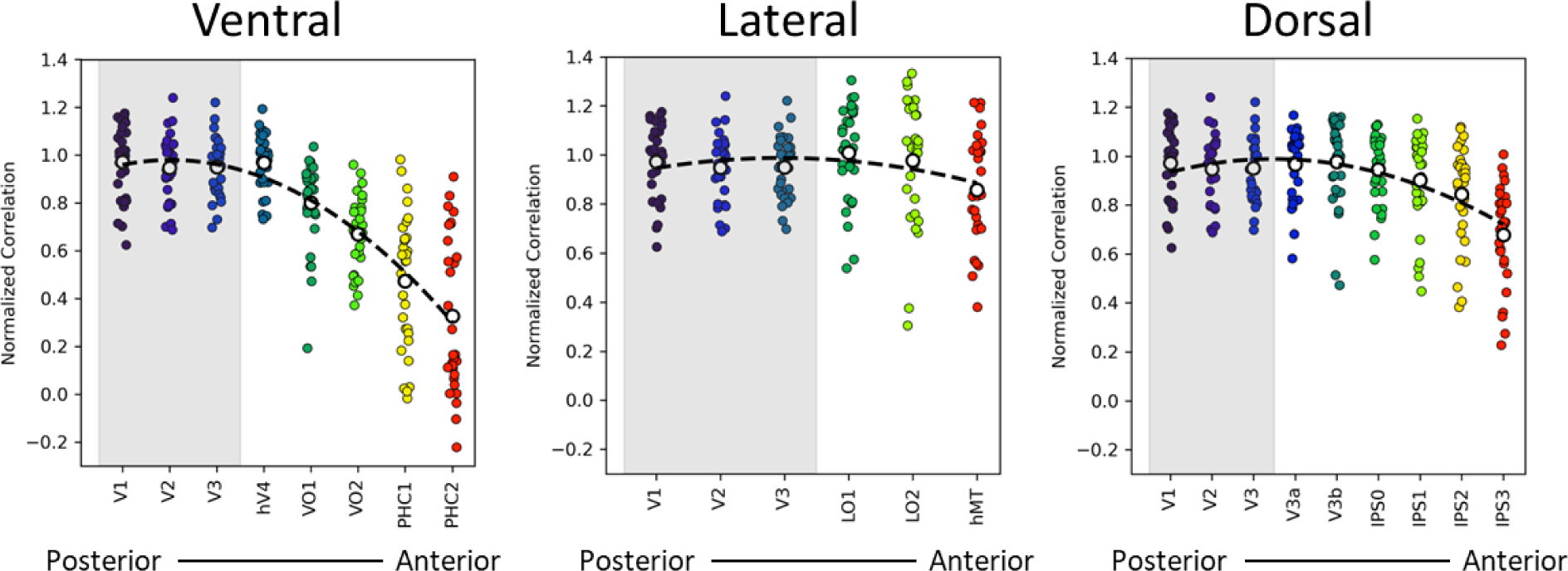
Differences in pulvinar-cortical connectivity correlations between neonates and adults within each visual pathways. Colored dots represent the normalized correlation between each individual neonate and the adult group map for each area. Colors correspond areal distinctions in Figure 3C. White dots represent the mean correlation between neonates and adults for each area. The dotted line indicates the fit of a two-feature polynomial. Normalized correlations were computed by scaling each individual neonate correlation to the adult noise-ceiling. The shaded region delineates occipital visual areas, which mark of the start of each pathway.

### Retinotopic connectivity within the pulvinar

Having characterized the organization and maturity of pulvino-cortical connections in neonates, we then examined the organization and specificity of pulvinar connectivity *within* an individual cortical area in neonates. In adults, connectivity between the pulvinar and visual cortex is retinotopically organized. To examine whether pulvino-cortical connectivity is retinotopically organized already at birth, we compared pulvinar connectivity patterns between dorsal and ventral portions of V1, corresponding to lower and upper visual field representations of visual space, respectively (Figure 6A). Computing a subtraction between the dorsal and ventral V1 tractography maps revealed spatially distinct connectivity within the pulvinar. Tracking with ventral V1 (blue; Figure 6B) was located relatively more ventral-lateral within the pulvinar and tracking with dorsal V1 (red) was located in an adjacent dorsal-medial portion. The organization of this connectivity corresponded to the retinotopic organization typically found within the adult ventral pulvinar (Arcaro et al., 2015). There was a significant rank-order correlation between polar angle representations defined in adults and the subtraction map between dorsal and ventral V1 probabilistic tractography in neonates (Spearman *rho* = 0.39, 95% CI [0.33, 0.47], *p* < .001) and in adults (Spearman *rho* = 0.43, 95% CI [0.39, 0.51], *p* < .001), with no significant difference between groups (as evidenced by overlapping confidence intervals). These findings indicate a retinotopic organization to pulvino-cortical connectivity in neonates, suggesting that the neonate pulvinar is already well-positioned to process visual information in a spatially specific manner.

**Figure 6.**
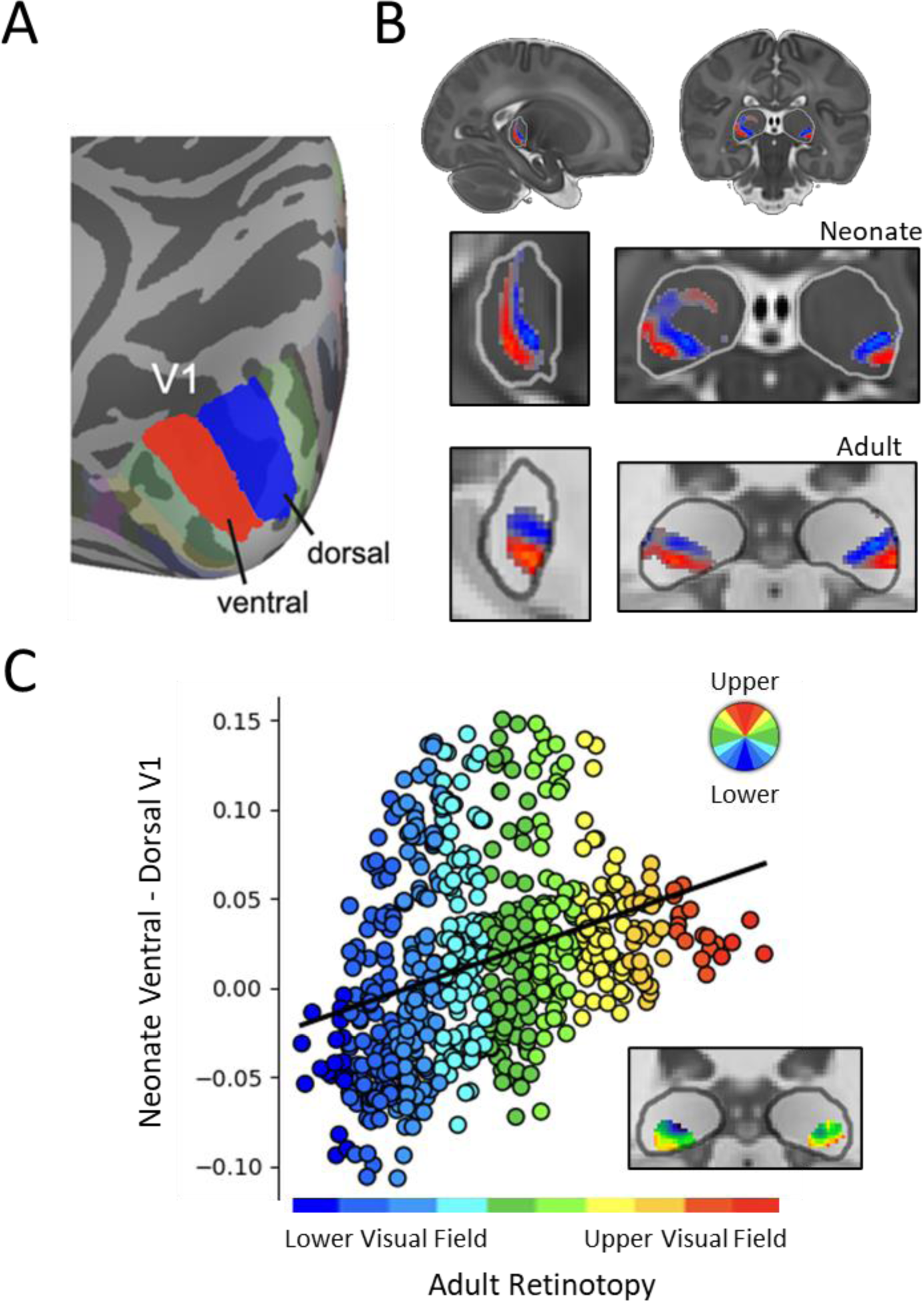
Retinotopic pulvino-cortical connectivity. (A) The anatomical location of dorsal and ventral V1, represented in blue and red respectively. (B) The coronal and sagittal view of the neonate (top) and adult (bottom) pulvino-cortical connectivity to ventral and dorsal V1 in the pulvinar. (C) The correlation between neonate V1 ventral-minus-dorsal connectivity and adult retinotopy. Values along the X-axis reflect the position of retinotopy values for adults, with blue representing values closer to the lower visual field and red reflecting values closer to the upper visual field. Values along the Y-axis reflect relative strength of dorsal (negative values) and ventral (positive values) V1 connectivity. The inset shows the retinotopic visual responses in the adult pulvinar.

## Discussion

Our investigation of pulvino-cortical anatomical pathways in newborn humans revealed extensive connections between the pulvinar and the entire visual system. The organization of these connections in neonates mirrored adult connectivity at multiple spatial scales of cortical organization, including functionally distinct cortical pathways, individual areas, and the intra-areal retinotopic organization. However, we also identified developmental differences in the maturity of connectivity patterns within the neonate pulvinar. Notably, pulvinar connectivity with areas of the ventral pathway were less adult-like than areas of occipital, lateral, and dorsal cortex, suggesting that pulvino-cortical connectivity along the ventral cortical pathway remains immature at birth. These findings highlight both the early establishment and ongoing maturation of pulvino-cortical connections prior to substantial experience and learning with the external world.

### Early scaffolding of pulvino-cortical connections

The large-scale organization of pulvino-cortical connections is already established at birth. Probabilistic tractography revealed the presence of white matter pathways between the neonate pulvinar and individual visual areas along the posterior thalamic radiation, aligning with prior work in adults (Arcaro et al., 2015; Le Gros Clark & Boggon, 1935; Leh et al., 2008). Each cortical area was interconnected with a distinct subregion within the pulvinar, with gradients of connectivity following the hierarchical organization of ventral, lateral, and dorsal cortical visual pathway. Consistent with adult tractography (Arcaro et al. 2015), connections with occipital areas were strongest in ventral-lateral portions of the pulvinar, with connectivity to higher-level extrastriate areas along the ventral pathway becoming stronger posteriorly. By contrast, connectivity to higher-level areas along the lateral and dorsal visual pathways became stronger dorsal-anteriorly. At a finer scale, intra-areal connectivity between V1 and the pulvinar reflected the retinotopic organization observed in the adults (Arcaro et al., 2015; Bender, 1982; Ungerleider et al., 1983), supporting the hypothesis that retinotopy is a common organizing principle the developing visual system (Arcaro & Livingstone, 2017). These findings demonstrate that pulvino-cortical connectivity at birth reflects multiple spatial scales of cortical organization, laying the foundation for complex and visual processing capabilities characteristic of the adult visual system.

### Differential development across pulvino-cortical pathways

Despite the presence of pulvinar connections throughout visual cortex in neonates, we observed developmental differences in the maturity of different visual clusters. Occipital areas V1-V3 showed adult-like connectivity patterns with the pulvinar, both in relation to the degree of anatomical overlap between neonate and adult white matter pathways and in the spatial profile of connections within the pulvinar itself. All neonate occipital areas showed patterns of connectivity at the adult noise ceiling that were distinct from other visual cluster patterns. These findings are consistent with extensive research showing that occipital cortex is one of the earliest cortical areas to mature (Bourne & Rosa, 2006; Ellis et al., 2021; Hubel & Wiesel, 1963; Le Vay et al., 1980). Our results extend these findings by showing that this maturation is reflected in the connectivity with the pulvinar.

Our results also contribute to the ongoing debate regarding the development of ventral and dorsal visual processing pathways (Ayzenberg & Behrmann, 2022; Braddick et al., 2003; Freud et al., 2019). While connections between the pulvinar and both ventral and dorsal visual cortex were identified, the dorsal pathway exhibited more adult-like connectivity patterns than the ventral pathway. Notably, ventral regions were the only network to exhibit correlations below the adult noise ceiling and their pattern of connectivity was less differentiable from the correlations with neighboring cortical areas (Figure 4). These findings are consistent with accumulating evidence of a protracted developmental trajectory for the ventral pathway relative to the dorsal pathway (Ayzenberg & Behrmann, 2024). The cytoarchitecture and functional properties of dorsal cortical regions are more mature than ventral regions in human children and juvenile monkeys (Bourne & Rosa, 2006; Ciesielski et al., 2019; Distler et al., 1996). Moreover, the organization of the ventral pathway is more shaped by postnatal experience, such that the functional profile of the ventral cortex is highly dependent an individual’s visual diet, such as seeing faces or reading text (Arcaro et al., 2019; Arcaro et al., 2017; Dehaene et al., 2010). The ventral pathway has a longer developmental window of plasticity than the dorsal pathway and is better able to reorganize following extensive cortical damage (Ahmad et al., 2022; Ayzenberg et al., 2023). One possibility, based on our results, is that the differential development of ventral and dorsal pathways is rooted in the initial immaturity of their connections with the pulvinar.

The lateral cortical pathway showed relatively mature connectivity with the pulvinar, with the correlations between neonates and adults reaching the noise ceiling and surpassing those of the ventral pathway (Figure 4_. However, some evidence of immaturity was observed in more anterior regions of the lateral pathway. Specifically, area MT showed less specificity in its spatial correlations and its correlation with adults was further from the noise ceiling than posterior lateral areas. This is particularly notable because an adult-like laminar organization of MT is present within the first postnatal month, indicating that this region matures earlier than other extrastriate visual regions (Bourne & Rosa, 2006; Mundinano et al., 2018). However, connectivity patterns between MT and the pulvinar change over postnatal development. Area MT receives greater input from the pulvinar than V1 in newborn marmosets, but, by adulthood, pulvinar connections to MT are substantially diminished and V1 becomes its primary source of input (Warner et al., 2012). Thus, the spatial profile of connectivity between the pulvinar and MT differs between neonates and older individuals. Our findings are consistent with these data, showing the early formation of connectivity between MT and the pulvinar, and that the connectivity patterns change between neonates and adult humans.

### Role of the pulvinar in visual development

Researchers have long suggested that early developing perceptual abilities are supported by subcortical, rather than cortical, structures (Blumberg & Adolph, 2023; Braddick & Atkinson, 2011; Johnson, 2005). For instance, newborn infants exhibit a motion perception asymmetry bias, such that they are particularly sensitive to objects moving in a temporal-to-nasal direction (Atkinson, 1979; Wattam-Bell, 2003), a bias consistent with the visual sensitivities of subcortical regions (Hoffmann, 1986). Furthermore, children with extensive resections to cortex, but intact subcortex, in one hemisphere (i.e., hemispherectomy) still show sensitivity to temporal-to-nasal motion (Morrone et al., 1999). Further, visually-evoked potential studies have shown that perception of object motion elicits cortical responses in infants older 6-weeks, but not earlier (Birch et al., 1985).

Other work has shown that newborn infants preferentially attend to face-like configurations (two dots above a third) before they have any visual experience with faces (Johnson et al., 1991; Reid et al., 2017; Sugita, 2008) or before they exhibit face-selective cortical responses (Arcaro et al., 2017). As with motion, there is a temporal-nasal asymmetry in neonates’ visual preferences for face-like stimuli (Simion et al., 1998). Moreover, neurons with sensitivity to face-like distribution of visual information are found in subcortical regions of non-human primates (Carlei et al., 2017; Hafed & Chen, 2016; Tsurumi et al., 2023; Yu et al., 2024), and, interestingly, a preference for face-like configurations is present even in evolutionarily-distant animals, such as turtles (Versace et al., 2020), suggesting that evolutionarily-preserved subcortical structures may support these abilities (Behrmann & Avidan, 2022). These findings are consistent with the two-process theory of face development: CONSPEC, an innate and evolutionarily old subcortical system that supports the initial detection of conspecific faces, and CONLERN a cortical system that develops as animals are exposed to the diagnostic features of individual faces (Morton & Johnson, 1991). Although more direct study with human infants is needed, the pulvinar may be well suited to support these various functions given that it is early developing (Shatz & Rakic, 1981), broadly connected with visual cortex (Arcaro et al., 2015; Kaas & Lyon, 2007; Shipp, 2003), and is known to contribute to these functions in adults (Arend et al., 2008; Eradath et al., 2021; Kagan et al., 2021; Wilke et al., 2010).

Finally, the pulvinar may play a central role in predictive error-driven learning (Cortes et al., 2023), also referred to as predictive-coding, thereby helping infants initially form a model of the world. Predictive learning is the idea that high-level regions of the brain continuously generate predictions about the future state of the world and these predictions are compared with bottom-up input about the actual current state (Friston, 2005). While higher association cortices are often the focus of predictive learning, the pulvinar is uniquely suited to generate error-driven signals in visual learning. This capability arises from its integration of strong bottom-up input from early visual cortex and modulatory top-down signals from higher-order visual areas (Bickford, 2016; Sherman & Guillery, 1998). Consistent with this hypothesis, the pulvinar shows earlier decoding of a stimulus embedded in noise than primary sensory areas (Barczak et al., 2018). Other work has shown that signals in the pulvinar reflected monkey’s confidence judgments during a motion discrimination task, and, indeed, temporary inactivation of the pulvinar led monkeys to opt-out of responding more often (Komura et al., 2013). Finally, biologically-plausible simulations of pulvino-cortical dynamics revealed that the pulvinar is both capable of implementing error-driven learning by integrating signal from occipital, ventral, and dorsal cortices, and that this learning leads to the development of invariant object recognition abilities (O’Reilly et al., 2021).

Given its disproportionately large size relative to other subcortical structures and its widespread connectivity to cortex, the pulvinar was once described as the “terra incognita” of the thalamus (Walker, 1966). Since then, researchers have uncovered the pulvinar’s various roles in visual processing and indicated it is crucial for the initial development of the visual system. Our results build on these findings by providing a detailed description of the extent and maturity of pulvino-cortical connections in human neonates. These data provide important constraints on theories of brain development and perceptual functioning in infancy.

## Methods

### Participants

Thirty neonatal scans ranging from 34.7 to 43.7 (*M* = 40.3) gestational weeks were randomly selected from the Developing Human Connectome Project (dHCP) dataset (Edwards et al., 2022). Data from a previously published study (Arcaro et al., 2015) on seventeen adults were re-analyzed for comparisons between age groups.

### Data acquisition & initial preprocessing

Neonatal MRI data were collected in a 3T Philips Achieva scanner with a 32-channel neonatal head coil and 80 mT/m gradient (Bastiani et al., 2019; Hughes et al., 2017; Makropoulos, Counsell, et al., 2018).

Diffusion data were collected with a multi-shell HARDI protocol (1.5 x 1.5 x 3mm with a 1.5mm slice overlap) using 4 different phase encoding directions at 3 b-value shells (400, 1000, and 2600 s/mm^2^) with different numbers of orientations per b-value shell (64, 88, and 128, respectively). T1-weighted and T2-weighted contrast anatomical images were collected using Turbo Spin Echo (TSE) and Inversion Recovery TSE sequences, respectively (0.8 x 0.8 x 1.6 mm resolution upsampled to 0.5 mm^3^ after image reconstruction). All scans were acquired without sedation. Initial data preprocessing including distortion correction (susceptibility, eddy currents, and motion) and segmentation and generation of individual subject cortical surfaces (Makropoulos, Robinson, et al., 2018) was provided by the dHCP group. Full details on data acquisition and initial preprocessing are reported for diffusion (Bastiani et al., 2019), anatomical imaging, and cortical surface reconstruction (Makropoulos, Robinson, et al., 2018).

### Data analysis

Analysis of Functional NeuroImages (AFNI; RRID:nif-0000-00259; Cox, 1996), SUMA (Saad and Reynolds, 2012), FreeSurfer (FreeSurfer, RRID:nif-0000-00304) (Dale et al., 1999; Fischl, 2012)(Dale et al. 1999; Fischl et al. 1999), FSL (FSL, RRID:birnlex_2067; Smith et al., 2004), Advanced Normalization Tools (ANTs; Avants et al. 2011), MATLAB (MATLAB, RRID:nlx_153890), and the NiLearn (Abraham et al., 2014)(RRID:SCR_001362), NumPy (RRID:SCR_008633), and Pandas (RRID:SCR_018214) packages for Python (RRID:SCR_008394) were used for additional data processing.

### Identification of cortical and subcortical visual areas

Analyses were performed between cortical retinotopic maps and the pulvinar. An adult probabilistic atlas of retinotopic maps (Wang et al., 2015) was used to identify putative visual areas in individual neonates. To align the adult maps to neonates, each neonate’s cortical surface was registered to an adult cortical surface template (FreeSurfer’s fsaverage). The registration process aimed to minimize misalignment by optimizing registration of five sulcal landmarks that span retinotopic maps (Supplemental Figure 1): 1) calcarine sulcus, 2) collateral sulcus, 3) lateral occipital sulcus, 4) parietal occipital sulcus, 5) frontal sulcus. These sulci were manually identified on each neonate’s cortical surface. For adults, sulci were identified using the DKT40 and Destrieux cortical surface atlases (Destrieux et al., 2010). Following registration, the adult probabilistic retinotopic maps were projected onto each neonate’s cortical surface. The accuracy of cortical surface registration was assessed by visually inspecting the localization of adult visual areas on the neonate cortical surface and confirming that the relationship between individual retinotopic maps and specific anatomical landmarks typically observed in adults was present in neonates. Specifically, we ensured that (1) the border between the dorsal and ventral components of V1 fell within the fundus of the calcarine sulcus (Arcaro et al., 2022), (2) PHC1-2 fell in the posterior portion of the collateral sulcus (Arcaro et al., 2009), (3) IPS0-3 fell along the superior bank of the intraparietal sulcus (Swisher et al., 2007), (4) V3A and V3B fell in the parieto-occipital sulcus (Press et al., 2001), and (5) LO1/2 fell within the lateral occipital sulcus (Larsson & Heeger, 2006). In total, 17 retinotopic areas were selected from the adult probabilistic atlas: V1, V2, V3, hV4, VO1, VO2, PHC1, PHC2, LO1, LO2, TO1(MT), V3A, V3B, IPS0, IPS1, IPS2, and IPS3 (Supplemental Figure 2). The maximum probability map for each retinotopic area was then projected from each neonate cortical surface into their high-resolution volumetric anatomical image and resampled to the resolution of the diffusion data for subsequent probabilistic tracking analysis. Note, that although TO1 is defined based on retinotopy, it is assumed to be functionally comparable to motion-selective area MT (Wang et al., 2015). We refer to this area as ‘MT’ so as to better connect our results with the existing thalamocortical literature (Dumbrava et al., 2001; Merabet et al., 1998; Warner et al., 2012). We excluded IPS4&5, SPL1, FEF, and TO2/MST from the analysis because the size of these areas in the group-averaged probabilistic atlas was outside the distribution of size measurements from individual participants.

To examine the large-scale organization of pulvino-cortical connectivity, retinotopic visual areas were grouped based on their anatomical localization within posterior occipital (V1-V3), ventral occipito-temporal (hV4, VO1-VO2, PHC1-PHC2), lateral occipito-temporal (LO1-LO2, MT), and dorsal extrastriate-parietal (V3A-V3B, IPS0-IPS3) cortex. These groupings were motivated by classic work that separates visual processing into distinct visual pathways (Kravitz et al., 2011; Kravitz et al., 2013; Milner & Goodale, 1993; Pitcher & Ungerleider, 2021), and recent work in adults that empirically clustered these retinotopic maps into four networks on the basis of resting state cortico-cortical correlation patterns (Haak & Beckmann, 2018). Here, we refer to these grouping of visual areas as ‘clusters’ based on their anatomical positioning. Variations in the assignment of individual retinotopic areas at the borders of these clusters (e.g., assigning hV4 to occipital vs. ventral cluster) did not qualitatively change the results.

The pulvinar and surrounding white matter were identified based on differences in the image intensities between grey and white matter (Arcaro et al., 2015). To ensure consistency across subjects, a region of interest (ROI) mask of the pulvinar was identified on the 40-week gestation template and then projected to each neonate’s native diffusion volumetric images using registration transformations provided by the dHCP group. For the initial probabilistic tracking, we ensured that the entire pulvinar was included in each subject by padding the pulvinar ROI ∼3mm on all sides. However, comparisons between neonates and adults were conducted using an ROI mask restricted to the pulvinar from the Morel Atlas (Supplemental Figure 2; Niemann et al., 2000).

### Probabilistic diffusion analyses

Probabilistic diffusion tractography was performed on Eddy-corrected data provided by the dHCP group. Crossing fibers were modeled with Bayesian Estimation of Diffusion Parameters Obtained using Sampling Technique (BEDPOSTX; Jbabdi et al., 2012) using the default dHCP parameters (3 fibers, 3000 burnin, an axially symmetric “zeppelins” model, Rmean = 0.3; Bastiani et al., 2019). Probabilistic tracking with crossing fibers (PROBTRACKX; Behrens et al., 2007) was then performed with 10000 samples, curvature threshold at 0.2, maximum number of steps of 2000, and step length of 0.5 mm. This diffusion tractography approach creates sample streamline trajectories starting from assigned seed regions and iterates the steps based on the voxel-wise orientation, as previously obtained from BEDPOSTX, until it reaches the criteria for termination. All probabilistic tracking was performed in each neonate’s native diffusion space within the left and right hemispheres separately.

### Exclusion criteria for tractography analyses

Exclusion masks were used to anatomically constrain streamline trajectories to proceed along white matter pathways that interconnect the thalamus and visual cortex in adults. To restrict tracking to the ipsilateral hemisphere, the contralateral hemisphere was included in each exclusion mask. To avoid streamlines crossing the ventricles, portions of the lateral ventricle surrounding the pulvinar were included in the exclusion mask. A single coronal slice, several millimeters anterior to the pulvinar, was included in the exclusion mask to focus all tracking posterior to the pulvinar. All exclusion masks were generated on the 40-week gestation template, projected to each neonate’s native diffusion space, and manually inspected for accuracy. Streamlines that entered the exclusion mask were discarded.

### Identification of pulvino-cortical white matter pathways in neonates

An initial probabilistic tractography analysis identified the thalamic radiations linking the visual cortex and the pulvinar in each neonate. Symmetric tracking was performed between the pulvinar and each cortical area such that the connectivity distribution from all voxels in the pulvinar that pass through the cortical area is summed with the connectivity distribution from all voxels in the retinotopic area that pass through the pulvinar. To assess tracking consistency across neonates for each cortical area, the streamline density maps (fdt_paths) were projected to the 40-week neonate template and then normalized by dividing the value of each voxel by the maximum value across all voxels for that tracking. This resulted in values ranging between 0 and 1 where a value of 1 indicated that the voxel had the highest streamline value while 0 indicated no streamlines were detected. Each normalized map then underwent thresholding and binarization, ensuring that only voxels encompassing a minimum 10% of all streamlines from the probabilistic tracking were included. This step eliminated low-probability tracks that extend beyond the thalamic radiation. Individual subject binarized maps were then averaged to produce group overlap maps where each voxel’s value corresponds to the number of neonates where above threshold tracking was identified. An additional group average was computed on the non-threshold, normalized data to visualize which voxels consistently had the most tracking in individual neonates. See Arcaro et al. 2015 for comparable analysis in adults.

The resulting neonatal thalamocortical tracks were compared to previously published data in adults (Arcaro et al., 2015) by computing the degree to which neonate pathways for each cortical area overlapped with the same pathways in adults. Overlap was computed as the percentage of neonate voxels for a pathway that fell inside the corresponding pathway in adults. This measure was used instead of mutual overlap (i.e., Dice coefficients), because the adult tracking data only included tracking to each cortical region as a whole (i.e., occipital, ventral, lateral, dorsal cortices), not individual areas as in neonates. Therefore, this measure enabled assessing whether the tracking for each individual visual area in neonates fell within the corresponding broader tracking observed in adults.

### Cortical anatomical connectivity within neonate pulvinar

A second probabilistic tractography analysis was conducted to examine the cortical connectivity patterns within the pulvinar. Tracking was performed between all voxels within the pulvinar ROI and each cortical area. To ensure tracking along known white matter pathways, a waypoint mask was included, that restricted tracking to the thalamic radiation identified in the initial analysis. For each cortical area, a section of the group average thalamic radiation, extending across multiple coronal slices in between that area and the pulvinar, was manually identified on the 40-week template. This mask was then projected into each neonate’s native diffusion space and designated as the waypoint mask. Distance correction was applied to adjust for the inherent drop in the number of streamlines with increasing distance from the seed mask. To ensure that the distance correction did not artificially amplify distal connections, tracking was also performed for each cortical area without the distance correction.

For each cortical area, the tractography analysis yielded a count for every pulvinar voxel indicating the number of probabilistic streamlines connected to the cortical area. These counts are referred to as pulvino-cortical tractography maps. The resultant single-subject maps were then projected to the 40-week gestation template. To compensate for variations among neonates due to nuisance factors that can impact overall tracking quality, such as differences in size between ROIs, the presence of crossing fibers, and variations in signal-to-noise ratio, the data were normalized. This normalization involved scaling the data to a range of 0 to 1, where a value of 1 corresponded to the voxel(s) with the highest streamline count for that particular tracking, while voxels with a value of 0 had no streamlines. The normalized data were then averaged to derive group means for each of the 17 retinotopic cortical areas. See Arcaro et al. 2015 for comparable analysis in adults.

### Hierarchical connectivity gradients within the neonate pulvinar

The organization of tractography within the neonate pulvinar was assessed in relation to the hierarchical organization of adult visual cortical areas into ventral, lateral, and dorsal visual pathways (Haak & Beckmann, 2018; Van Essen et al., 1992). To this end, we computed tractography gradients extending from the most posterior area of the occipital cortex (i.e., V1) to the most anterior areas of each pathway. Broadly, this was accomplished by assigning each pulvinar voxel a value that corresponds to the visual area with the strongest connectivity to that voxel.

More specifically, each visual cortical area was first assigned an index value corresponding to its anatomical position in a pathway’s hierarchy relative to V1 (Ventral: V1 = 1, V2 = 2, V3 = 3, V4 = 4, VO1 = 5, VO2 = 6, PHC1 = 7, PHC2 = 8; Lateral: V1 = 1, V2 = 2, V3 = 3, LO1 = 4, LO2 = 5, MT = 6; Dorsal: V1 = 1, V2 = 2, V3 = 3, V3A = 4, V3B= 5, IPS0= 6, IPS1= 7, IPS2= 8, IPS3 = 9). Then we computed the strength of connectivity to each visual area relative to its neighbor in the hierarchy using a subtraction method. From each cortical area we subtracted the group averaged normalized streamline map of the area posterior to it. For V1, the most posterior area, we reversed this process and subtracted V2 from V1. This subtraction process resulted in a map for each area wherein positive voxel values correspond to stronger tracking for the current visual area being analyzed, whereas negative values correspond to stronger tracking to the posterior area neighboring it (except for V1, where negative values corresponded to V2). Then we extracted the positive values (corresponding to the current area being analyzed) and scaled them between 0 and 1. We add these values to the current area’s index, thereby converting pulvinar voxel values into scalars that represent the strength of their connectivity to a particular area. Finally, all maps are layered on top of one another to create a single map illustrating the gradient of connectivity to each area within a pathway. As an example, pulvinar voxel values between 2 and 3 are those whose connectivity is stronger with V2 than V1, whereas values between 3 and 4 are those whose connectivity is stronger to V3 than V2. Thus, the final map for each visual pathway is organized such that voxels with the lowest values correspond to the posterior areas of the pathway (e.g., V1, V2, V3), middle values correspond to intermediate areas (e.g., hV4, LO1, V3B), and larger values correspond to the most anterior areas (e.g., PHC2, MT, IPS3). Occipital areas (V1-V3) are included in each gradient map because they are the common starting point for each pathway and because they serve as a baseline of comparison for intermediate and anterior areas across pathways.

### Quantifying the maturity of pulvinar connectivity in neonates

To examine the maturity of pulvinar connectivity with each cortical area, the normalized streamline maps from each neonate were compared to previously published adult data (Arcaro et al., 2015). To this end, we conducted Pearson correlations between the spatial pattern for individual neonate maps with the adult group average pulvino-cortical tractography maps (e.g., individual neonate V1 vs. group adult V1). Similarity across age groups was examined at two levels-of-analysis: individual area and anatomical cluster of areas. In the cluster-level analysis, correlations between neonate and adult areas were averaged within-cluster (e.g., average of all occipital-to-occipital correlations) and compared to the average correlation between clusters (e.g., occipital-to-lateral correlations). In the area-level analysis, we tested whether the similarity between neonate and adult pulvinar tractography maps for an individual cortical area was greater than the similarity to other areas within-the same cluster (e.g., neonate V1 to adult V1 vs. neonate V1 to adult V2). Finally, to estimate how adult-like each tractography map was, we computed a noise-ceilings for each cortical area in adults using a leave-one-out cross validation procedure. For each adult participant and area, the tractography map was correlated with the group-averaged tractography map of the remaining adult participants, and the resulting correlations from all permutations were averaged together. This provided an estimate of the upper-bound correlation that could be found when correlating neonates to adults.

### Retinotopically-organized tractography

Tractography between the neonate pulvinar and V1 was compared to previously published retinotopic mapping within the adult pulvinar. The V1 retinotopic map was divided into dorsal and ventral portions, representing the lower and upper visual fields, respectively. Probabilistic tracking analyses were conducted to identify pulvinar voxels with the highest probability of connections to each portion of the V1 map in each neonate. Normalized group averaged pulvino-cortical connectivity maps were calculated for the ventral and dorsal portions of V1 and projected onto the adult MNI template. The dorsal map was subtracted from the ventral map, resulting in a new pulvinar map illustrating the relative difference in tracking with upper and lower visual field portions of V1. This difference map was then correlated with an adult group average map of polar angle representations in the ventral pulvinar.

## Supplemental Materials

**Supplemental Figure 1.**
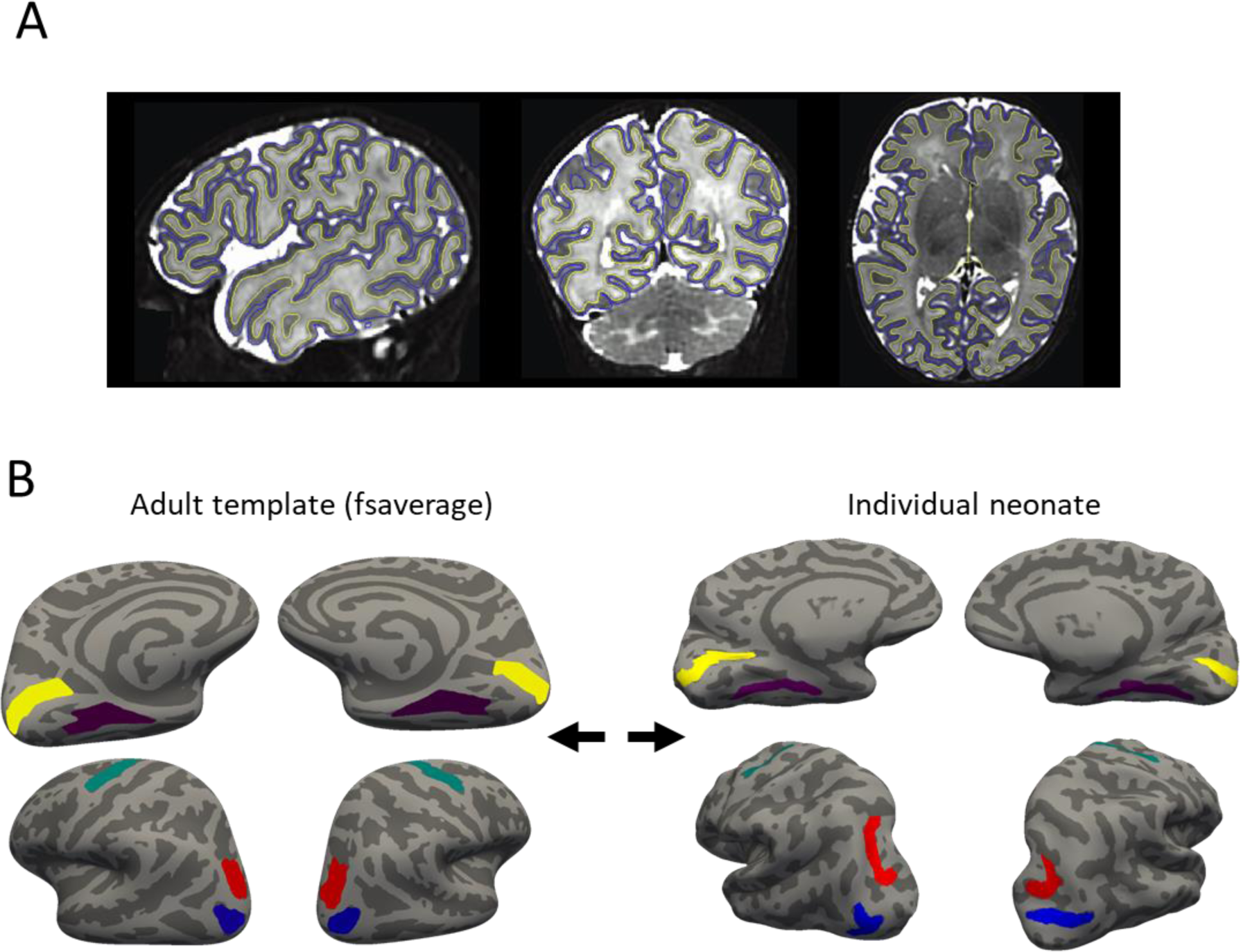
(A) An example of a cortical segmentation used to generate the surface for an individual neonate. (B) An illustration of the sulcal registration procedure. Registration between the adult template and each individual neonate was designed to minimize error between the (yellow) calcarine sulcus, (purple) collateral sulcus, (blue) lateral occipital sulcus, (red) parietal occipital sulcus, and (teal) frontal sulcus.

**Supplemental Figure 2.**
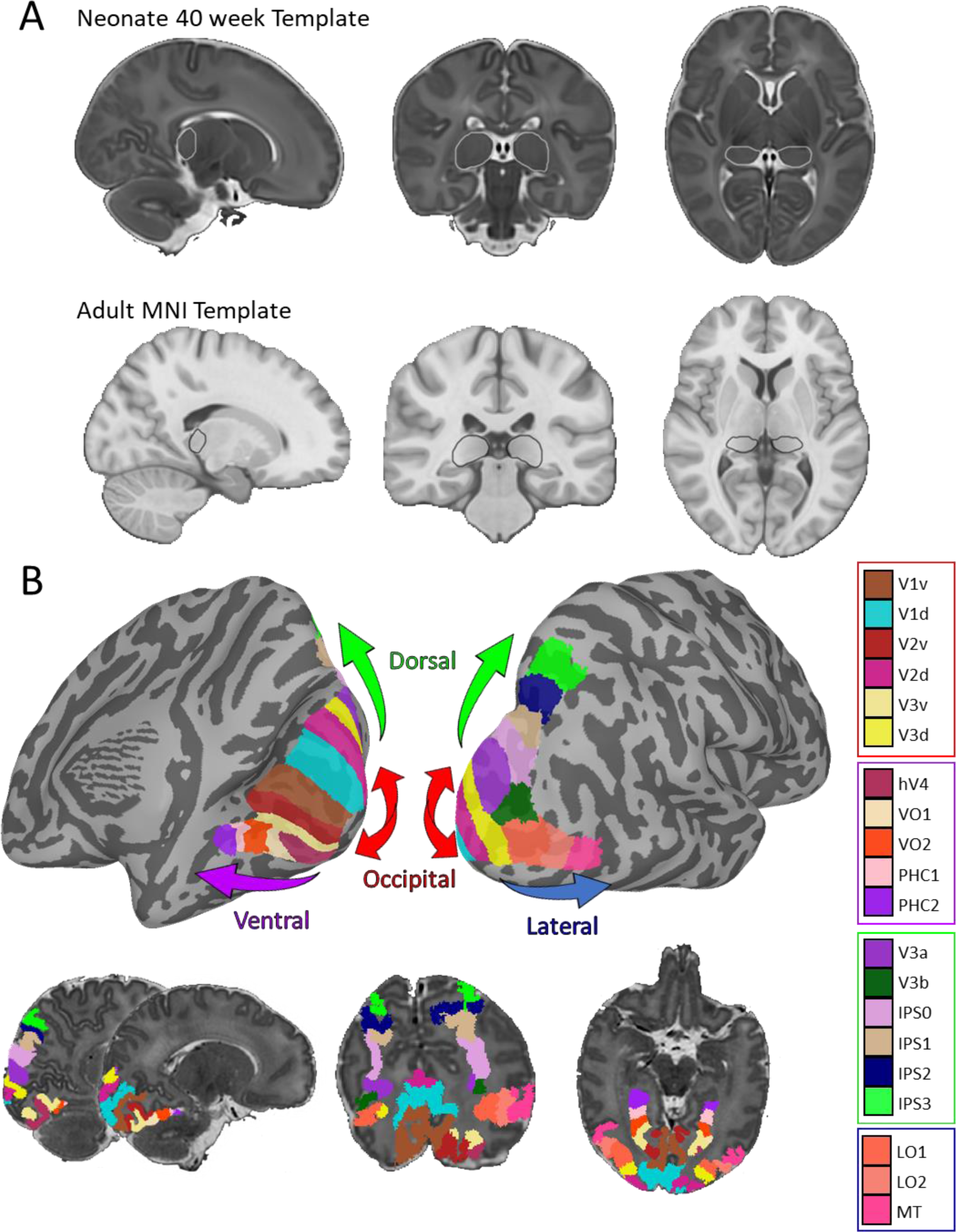
Pulvinar and retinotopic visual maps. (A) Morel pulvinar ROI projected onto (top) neonate 40-week template and (bottom) adult MNI template. (B) Retinotopically defined visual areas projected onto the same neonate’s brain in (top) surface and (bottom) volumetric space.

**Supplemental Figure 3.**
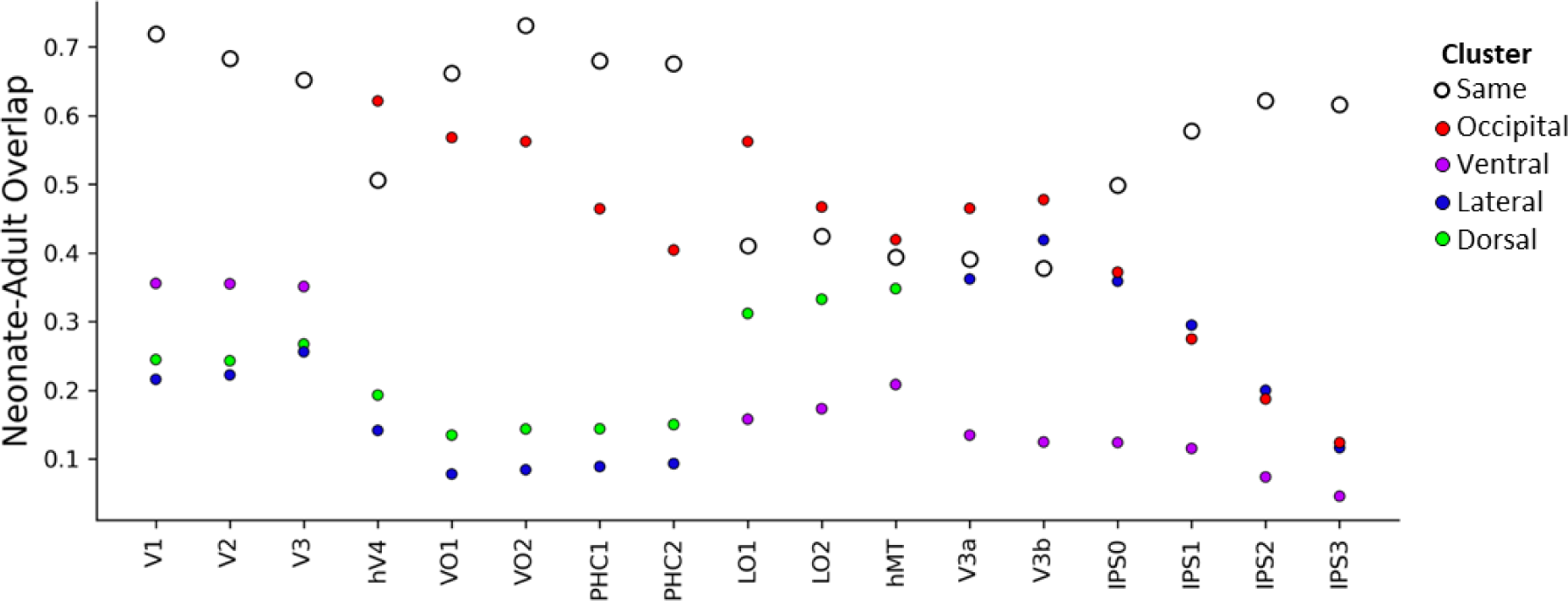
The degree of anatomical overlap between white matter pathways connecting the pulvinar and each cortical visual area in neonates with adults. White dots indicate the overlap with the matched network in adults and colored dots indicate the overlap with other visual networks. Color coding of visual region matches conventions in Figure 1. Overall, individual visual areas in neonates showed adult-like specificity in the location of pulvino-cortical white matter pathways (Occipital areas: *ds* > 6.86; *ps* < .001; Ventral areas: *ds* > 2.07; *ps <* .001; Lateral areas: *ds* > 1.11, *s ps* < .001; Dorsal areas: *ds* > 0.66, *ps* < .003).

**Supplemental Figure 4.**
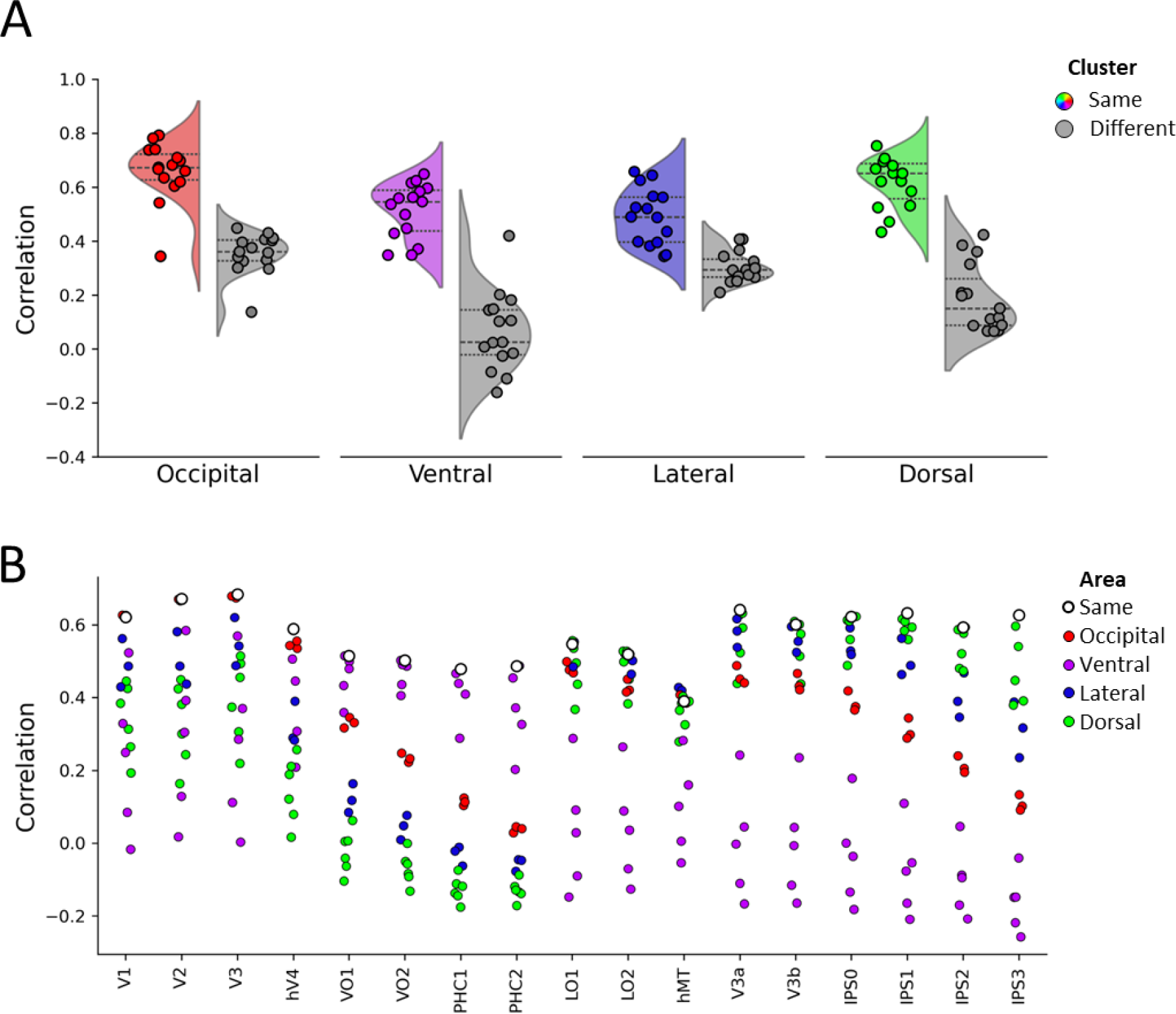
Connectivity map comparisons within the pulvinar between individual adult participants and the adult group maps. Adult data was analyzed using a leave-one-out procedure, wherein each individual adult’s connectivity map was correlated to a group-averaged map of the remaining adult participants. (A) Cluster-level similarity between adults for occipital, ventral, lateral, and dorsal visual areas. Colored distributions show the correlation between anatomically clustered visual areas in neonates with the same cluster in adults. Gray distributions show the correlation with different clusters. (B) Area-level similarity between adults for all visual areas measured. White dots indicate the correlation with the matched area and colored dots indicate the correlation with other visual areas. The other visual areas are color coded by anatomical cluster.

